# SORLA mediates endocytic uptake of proIAPP and protects against islet amyloid deposition

**DOI:** 10.1101/2022.04.21.488729

**Authors:** Alexis Z.L. Shih, Yi-Chun Chen, C. Bruce Verchere, Thomas E. Willnow

## Abstract

**Aims/ hypothesis:** Sorting-related receptor with type A repeats (SORLA) is a neuronal sorting receptor that prevents accumulation of amyloid-beta peptides, the main constituent of senile plaques in Alzheimer disease. Recent transcriptomic studies show that SORLA transcripts are also found in pancreatic islet beta cells, yet the role of SORLA in islets is unclear so far. Based on its protective role in reducing amyloid burden in the brain, we hypothesized that SORLA may have a similar function in the pancreas, regulating islet amyloid plaque formation from islet amyloid polypeptide (IAPP).

**Methods:** We generated human IAPP transgenic mice lacking SORLA (hIAPP:SORLA KO) to assess the consequences of receptor deficiency for islet histopathology and function *in vivo*. Using both primary islet cells and established cell lines, we further investigated the molecular mechanisms whereby SORLA controls the cellular metabolism and accumulation of IAPP.

**Results:** Loss of SORLA activity in hIAPP:SORLA KO resulted in a significant increase in islet amyloid deposits and associated islet cell death as compared to hIAPP:SORLA WT animals expressing the receptor. Aggravated islet amyloid deposition was observed in mice fed a normal chow diet, not requiring high-fat diet feeding typically needed to induce islet amyloidosis in mouse models. Further *in vitro* studies showed that SORLA binds to and mediates the endocytic uptake of proIAPP, but not mature IAPP, delivering the propeptide to an endolysosomal fate.

**Conclusions/interpretation:** SORLA functions as a clearance receptor specific for proIAPP, protecting against islet amyloid deposition and associated cell death caused by IAPP.

## Introduction

Islet amyloid polypeptide (IAPP or amylin) is a peptide hormone secreted together with insulin by pancreatic islet beta cells. IAPP is synthesized as a 67 amino acid precursor (proIAPP_1-67_) with two propeptide regions extended at its N- and C-terminus. Both proIAPP_1-67_ and its partially processed intermediate proIAPP_1-48_ can be released constitutively, while the 37 amino acid mature IAPP undergoes further processing and modifications in the secretory granules prior to glucose-stimulated secretion [1, 2]. IAPP contributes to the maintenance of energy homeostasis, in part through mediating satiety, delaying gastric emptying and modulating insulin secretion [3, 4]. However, human IAPP (hIAPP) is amyloidogenic and susceptible to aggregation into toxic oligomers and insoluble islet amyloid, a pathological feature of type 2 diabetes [5, 6]. It has been suggested that impaired proIAPP processing and overproduction of (pro)IAPP promote islet amyloid deposition [7, 8]. Still, the regulation of IAPP production and turnover, underpinning islet amyloid formation remains incompletely understood.

Similar to type 2 diabetes, accumulation of amyloid plaques is a pathological hallmark in Alzheimer disease (AD) [9]. The amyloidogenic agent in AD is amyloid-beta peptide (Aβ), a proteolytic product of the amyloid precursor protein (APP). Accumulation of Aβ in the brain is controlled by SORLA (sorting protein-related receptor containing LDLR class A repeats), a type I transmembrane receptor and major AD risk gene [10]. SORLA reduces overall Aβ burden through two mechanisms. It acts as an intracellular sorting receptor for APP, moving the precursor from endosomal compartments to the Golgi to prevent proteolytic breakdown to Aβ in endosomes [11, 12]. In addition, it sorts newly produced Aβ to lysosomes for catabolism, further reducing build-up of Aβ in the brain parenchyma [13, 14]. In line with a central role for SORLA in brain amyloidosis, genetic variants in *SORL1*, the gene encoding this receptor, have been associated with sporadic [15, 16] and familial [17] forms of AD.

While most studies have focused on the relevance of SORLA for amyloidogenic processes in neurons, recent single-cell RNA sequencing analyses of human and mouse islets uncovered SORLA transcripts also to be expressed in pancreatic islet beta cells [18–20]. Since SORLA is known to bind a broad range of ligands, including the amyloidogenic peptide Aβ [13, 21], we hypothesized that this receptor may have a role in beta cell physiology, specifically in regulating IAPP handling and islet amyloid formation. In this study, we tested if and how SORLA controls IAPP trafficking and processing, islet amyloid deposition, and glucose homeostasis. We examined the expression and subcellular localization of SORLA in islet beta cells and its interaction with pro- and mature forms of IAPP. Furthermore, we investigated the consequences of receptor deficiency for islet amyloid formation, beta cell function and overall metabolic characteristics in mice expressing hIAPP, an animal model of islet amyloid deposition.

## Results

### SORLA is expressed in islet beta cells

To validate existing transcriptome data on SORLA expression in islets on the protein level, we first examined SORLA expression by immunohistology in mouse and human pancreatic sections. In mouse islets, SORLA was mainly expressed in insulin-producing beta cells, but to some extent also in glucagon-producing alpha cells, somatostatin-producing delta cells and pancreatic polypeptide-producing PP cells (Fig.1a). Expression of SORLA was lost in islets of mice carrying a targeted disruption of *Sorl1*, hereinafter referred to as SORLA knockout (KO) (Fig.1b) [12]. In human islets, SORLA was predominately expressed in beta cells and only occasionally found in alpha cells (Fig.1c). Overall, these data document comparable expression patterns for SORLA in murine and human islets, consistent with prior transcriptome data [18–20].

**Fig.1.**
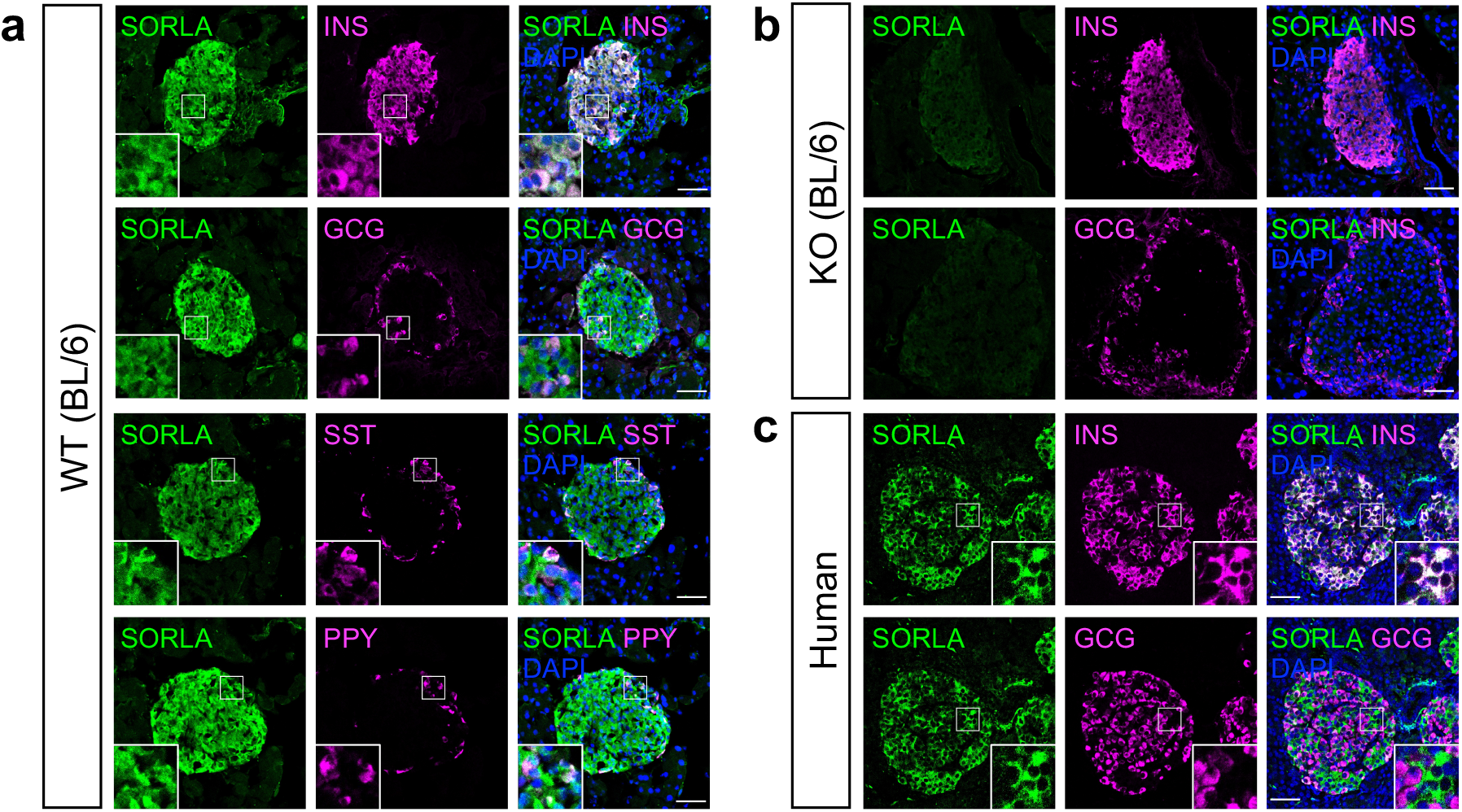
Expression of SORLA in murine and human pancreatic islets. (**a, b**) Immunofluorescence staining of pancreatic sections from (a) WT (BL/6) and (b) SORLA KO (BL/6) mice for SORLA (green), insulin (INS, magenta), as well as glucagon (GCG, magenta), somatostatin (SST, magenta), and pancreatic polypeptide (PPY, magenta). Nuclei were counterstained with DAPI (blue). Prominent expression of SORLA is seen in WT but not in SORLA KO islets. (**c**) Immunodetection of SORLA (green) and insulin or glucagon (magenta) on sections of human pancreatic biopsies from non-diabetic patients. Single and merged channel configurations are shown. The insets depict higher magnifications of the areas indicated by white boxes in the overview images. Scale bars, 50 μm.

### Loss of SORLA increases islet amyloid deposition

To study the impact of SORLA activity on islet amyloid formation, we crossed SORLA KO mice (on C57BL/6J background) with a transgenic line expressing hIAPP under the control of the rat insulin II promoter (on FVB/N background) [22, 23]. Male mice heterozygous for the hIAPP transgene and genetically deficient for *Sorl1* (hIAPP:SORLA KO), as well as hIAPP-expressing control animals (hIAPP:SORLA WT) were selected for analysis. We also generated non-transgenic control groups (SORLA WT, SORLA KO) to assess the impact of SORLA deficiency on glucose homeostasis in the absence of the hIAPP transgene. In addition to age, dietary stress imposed by high-fat diet (HFD) feeding contributes to islet amyloid formation [24]. Therefore, we performed our studies in mice fed a normal chow diet (ND) or a HFD (60% crude fat) for 6 months.

We first examined the effects of SORLA deficiency on islet amyloid deposition by staining pancreatic tissue sections from 7-month old hIAPP:SORLA WT and hIAPP:SORLA KO mice with thioflavin S (ThioS). Remarkably, a significant amount of islet amyloid was already evident in hIAPP:SORLA KO mice on ND (6.9±1.8% ThioS positive area of total islet area), while virtually none was observed in hIAPP:SORLA WT mice (0.56±0.16% ThioS positive area of total islet area) (Fig. 2). After HFD feeding, hIAPP:SORLA KO mice still tended to develop more islet amyloid (7.8±2.6% ThioS positive area of total islet area) compared to hIAPP:SORLA WT (2.9±0.8% ThioS positive area of total islet area), but this effect was not statistically significant (Fig. 2).

**Fig. 2.**
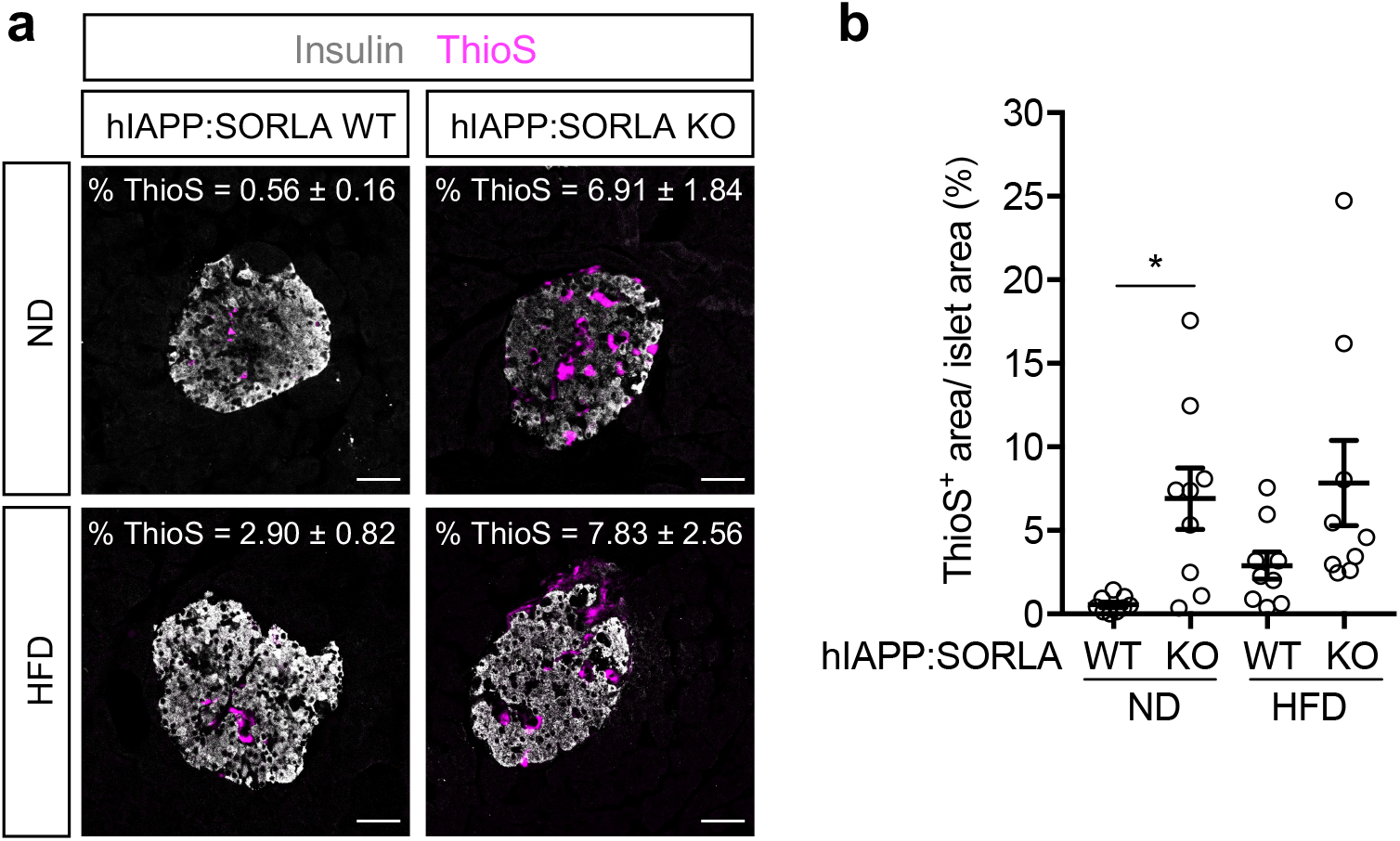
SORLA deficiency promotes islet amyloid deposition in hIAPP-expressing mice. (**a**) Amyloid deposition was evaluated on pancreatic sections from 33- to 35-weeks old mice of the indicated genotypes by staining with thioflavin S (ThioS, magenta) and immunostaining for insulin (white). Animals were fed a normal chow diet (ND) or a high-fat diet (HFD) for 6 months. Scale bars, 50 μm. (**b**) Quantification of ThioS^+^area per islet area on replicate pancreatic sections as exemplified in panel (a) (*n* = 9 mice per genotype, 20 – 30 islets per mouse). Data are shown as mean ± SEM. Statistical significance of differences was determined by two-way ANOVA with post hoc test. * *p* < 0.05.

Islet amyloid is known to promote islet cell death and impair beta cell function. Accordingly, we performed TUNEL staining to test whether the increased islet amyloid observed in SORLA-deficient mice correlated with a higher level of cell death. In line with an increased islet amyloid burden, the percentage of apoptotic cells per islet was significantly higher in hIAPP:SORLA KO mice as compared to hIAPP:SORLA WTs. Again, this defect was seen in mice fed a ND but not a HFD (Fig. 3). SORLA deficiency specifically impacted amyloid burden and cell survival but not overall islet morphology, as neither total islet area nor the proportions of beta cell or alpha cell per islet were significantly different between genotypes (Suppl. Fig.1).

**Fig. 3.**
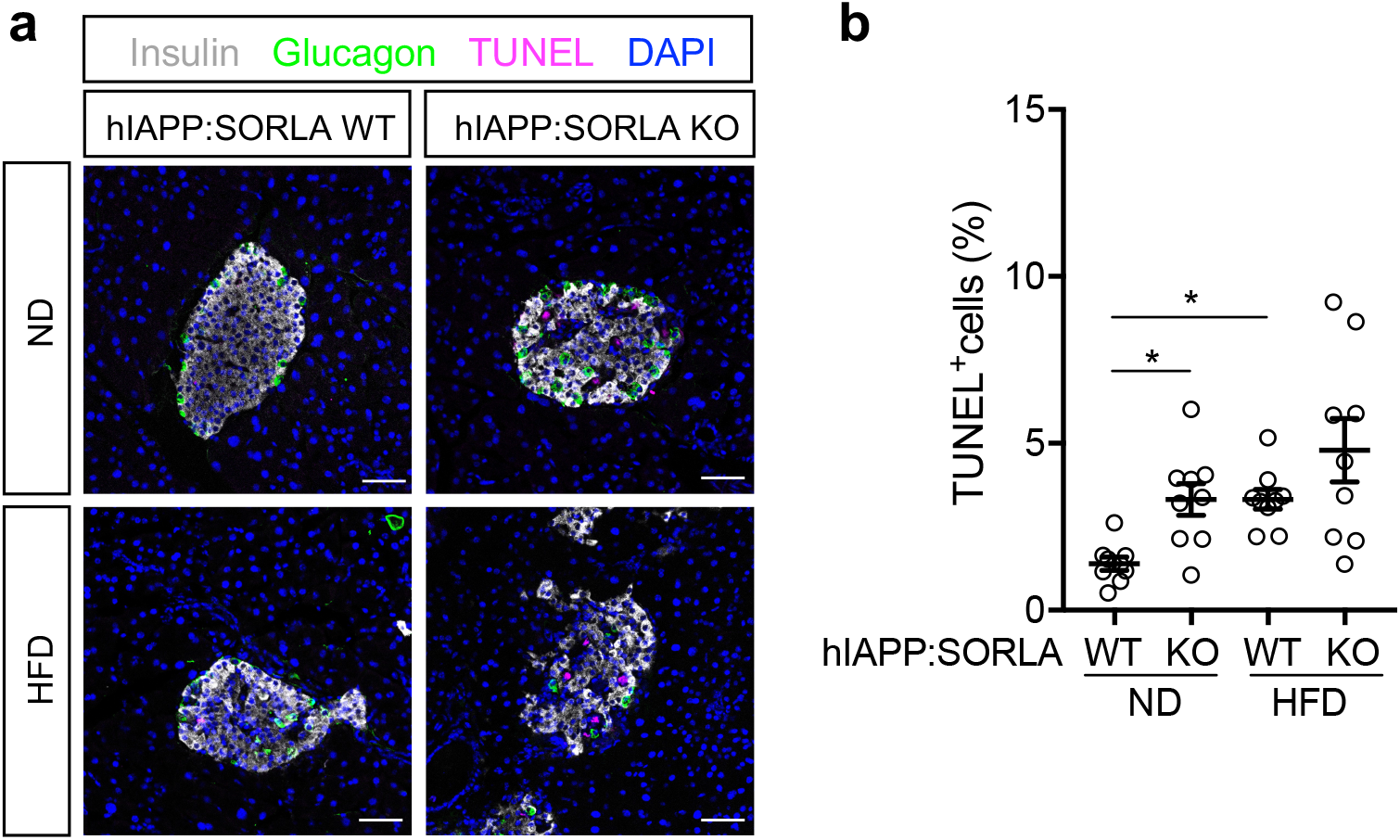
Enhanced islet amyloid deposition in SORLA-deficient mice correlates with increase islet cell death. (a) Apoptotic cell death was evaluated on pancreatic sections from 33- to 35-weeks old mice of the indicated genotypes by TUNEL staining (magenta). Sections were also immunostained for insulin (white) and glucagon (green). Nuclei were counterstained with DAPI (blue). Animals were fed a ND or HFD for 6 months. Scale bars, 50 μm. (**b**) Quantification of the percentage of TUNEL^+^ cell per islet on replicate pancreatic sections as exemplified in panel a (*n* = 9 mice per genotype, 20 – 30 islets per mouse). Data are shown as mean ± SEM. Statistical significance of differences was determined by two-way ANOVA with post hoc test. * *p* < 0.05.

### Glucose metabolism and beta cell function of hIAPP-expressing SORLA KO mice

In addition to histological assessments of islet amyloid burden at endpoint, we also monitored the metabolic consequences of SORLA deficiency in both hIAPP-transgenic and non-transgenic mice over time. ND-fed SORLA-deficient animals were slightly heavier than their WT littermates from around 26 weeks of age (Fig. 4a). However, SORLA deficiency did not impact fasting blood glucose levels, regardless of hIAPP transgene expression (Fig. 4b). Expression of the hIAPP transgene resulted in impaired glucose tolerance when compared with non-transgenic controls (Fig. 4c), while SORLA KO animals showed normal glucose tolerance compared to WT controls (Fig. 4c). Additionally, we determined the effects of SORLA on beta cell function by measuring glucose-stimulated insulin secretion in mice (Fig. 4d) and isolated islets (Fig. 4e). The fold change in insulin secretion upon intraperitoneal glucose administration was similar between SORLA genotypes (Fig 4d). Similarly, SORLA deficiency did not impact glucose- or KCl-stimulated insulin secretion in perifusion experiments of islets from hIAPP-expressing mice (Fig. 4e).

**Fig. 4.**
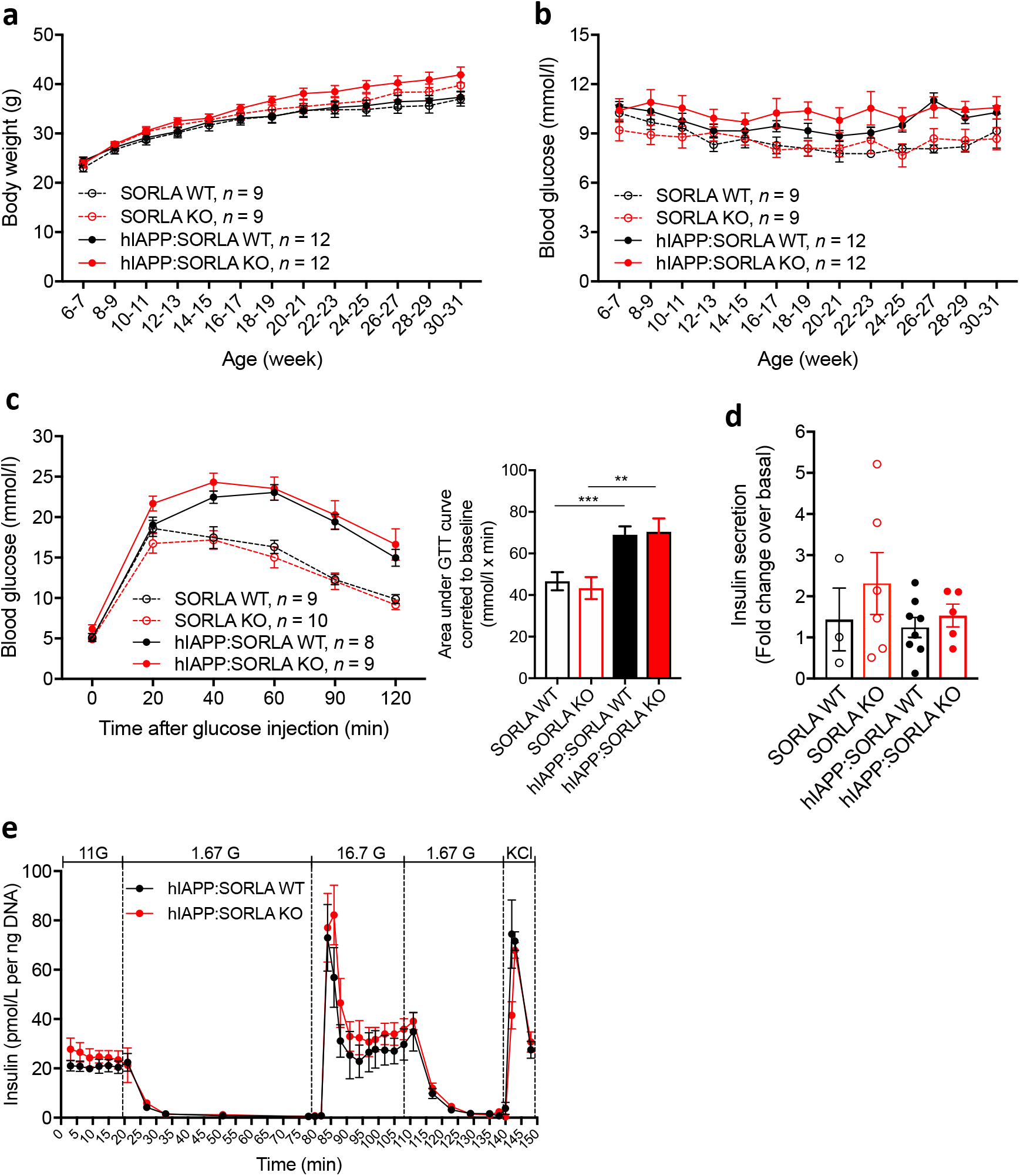
Metabolic characterization and beta cell function of hIAPP-expressing wildtype and SORLA-deficient mice on a normal chow diet. (**a-b**) Bi-weekly analysis of mice of the indicated genotypes for (a) body weight and (b) 6-h fasting blood glucose levels. **(c)** Glucose tolerance test (GTT) performed in 30- to 32-weeks old mice after a 16 h fast by intraperitoneal injection of 2 g/kg body weight of glucose. Response to glucose clearance was quantified based on the area under the GTT curves corrected to baseline. **(d)** Glucose-stimulated insulin secretion (GSIS) was performed in 31- to 33-weeks old mice of the indicated genotypes (SORLA WT, *n* = 3; SORLA KO, *n* = 6; hIAPP:SORLA WT, *n* = 8; hIAPP:SORLA KO, *n* = 5). A glucose dose of 2 g/kg body was administered intraperitoneally after a 16 h fast. Data are represented as fold change in insulin secretion at 30 minutes post-glucose injection over basal level. **(e)** Dynamic GSIS was tested on perifused islets from 31- to 33-weeks old hIAPP-expressing SORLA WT or KO mice (*n* =3 mice per genotype, with technical duplicates). Data are shown as mean ± SEM. Statistical significance of differences in (c) was determined by unpaired Student t-test. ** *p* < 0.01, *** *p* < 0.001.

To further determine if SORLA promotes the development of diabetes (dependent or independent of hIAPP), we challenged mice with HFD for 6 months starting at 4 weeks of age. SORLA-deficient mice grew heavier on HFD compared to their WT littermates (Suppl. Fig. 2a). Yet, similar to ND-fed mice, SORLA-deficient mice on HFD had fasting blood glucose levels (Suppl. Fig. 2b) and glucose tolerance (Suppl. Fig. 2c) comparable to their WT littermates.

Taken together, our *in vivo* data demonstrated that loss of SORLA promotes islet amyloid burden and islet cell loss without overt impact on blood glucose homeostasis. This defect was observed in normal chow-fed mice and did not require any dietary stressor. These findings suggest a possible role for SORLA in protecting the pancreas from early stages of islet pathology preceding signs of diabetes.

### Loss of SORLA does not alter proIAPP processing

Next, we aimed at elucidating the molecular mechanism whereby SORLA controls IAPP handling and islet amyloid deposition. Since impaired processing of proIAPP has been implicated in islet amyloid formation [1, 7, 25], we initially tested whether SORLA may function as a sorting receptor regulating IAPP maturation in islet beta cells, similar to its function in sorting APP in neurons [12, 14]. Accordingly, we measured fasting plasma levels of prohIAPP_1-48_ and mature hIAPP in our mouse models by ELISA and calculated the ratio of pro- to mature hIAPP as an indicator of processing efficiency. Both hIAPP-expressing SORLA WT and KO mice had similar levels of mature hIAPP, prohIAPP_1-48_, and ratio of pro- to mature hIAPP whether fed a ND (Suppl. Fig. 3a-c) or HFD (Suppl. Fig. 3d-f). To further validate these *in vivo* results, we measured levels of mature hIAPP and prohIAPP_1-48_ secreted from isolated islets in standard glucose culture (11 mmol/l), and following low (1.67 mmol/l) or high glucose (16.7 mmol/l) conditions (Suppl. Fig. 3g). No significant differences in the amounts or ratios of secreted mature hIAPP and prohIAPP_1-48_, (Suppl. Fig. 3h - j), nor in total islet contents of mature and prohIAPP_1-48_ (Suppl. Fig. 3k, l), were noted in SORLA genotypes across all conditions. These data suggest that SORLA does not directly or indirectly impact proIAPP processing.

### SORLA is a receptor for IAPP

Prior studies in various cell types have shown that SORLA mainly localizes to the Golgi, cell surface, and endosomes [26], in line with its function in sorting cargo between plasma membrane, secretory and endocytic compartments. We tested if SORLA localizes to similar subcellular compartments in beta cells by co-immunostaining for SORLA and organelle markers in dispersed islet cells from WT (BL/6) mice (Fig. 5a). Quantification of double-immunostained cells using Manders’ colocalization coefficient showed that SORLA is most highly colocalized with secretory granule markers (insulin and IAPP); moderately with early (Rab4), late (Rab9), and recycling endosomes (Rab11); and to a lesser extent with the *trans-Golgi* network (STX6) (Fig. 5b).

**Fig. 5.**
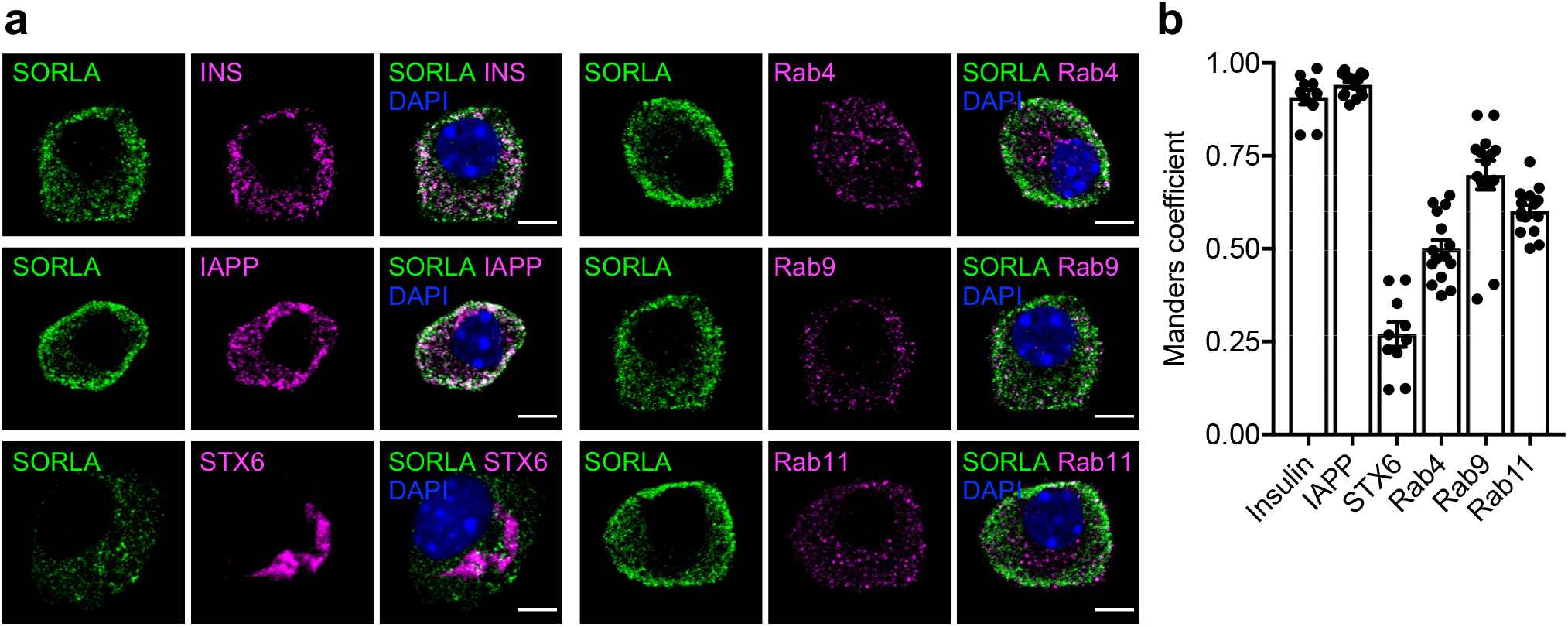
SORLA colocalizes with IAPP to the endocytic compartment of islet beta cells. (a) Dispersed islet cells from WT(BL/6) mice were immunostained for SORLA (green) and the following cell compartment markers (magenta): insulin or IAPP (secretory vesicles), syntaxin-6, STX6 *(trans-Golgi)*, Rab4 (early endosomes), Rab9 (late endosomes), or Rab11 (recycling endosomes). Nuclei were counterstained with DAPI. Single as well as merged channel configurations are shown. Scale bars, 5 μm. (b) Degree of colocalization between SORLA and each compartment marker was quantified by Manders’ colocalization coefficient (*n* =10-15 cells). Data are shown as mean ± SEM.

To examine whether SORLA is able to act as a receptor for pro- or mature forms of IAPP, we tested their binding interactions using microscale thermophoresis (MST). In this experiment, we used a His-tagged version of the human SORLA ectodomain (Fig. 6a), recombinantly expressed and purified from HEK293-EBNA cells as previously described [12]. Ligand binding of the ectodomain was tested using commercially synthesized forms of mouse N-terminally extended proIAPP (proIAPP_1-70_), C-terminally extended proIAPP (proIAPP_1-51_) or mature IAPP peptides (Fig. 6b). The non-aggregating mouse IAPP peptides were chosen as they exhibit better solubility and stability as monomers *in vitro* compared to the aggregating human form [27]. Binding interactions were detected by comparing multiple measurements over a temperature gradient, with a constant concentration of fluorescently-labeled SORLA ectodomain and a serial titration of unlabeled IAPP peptides. Using the K_*d*_ model of fit evaluated by Affinity Analysis software, the SORLA ectodomain was shown to interact with all three forms of IAPP (Fig. 6c). However, binding was substantially stronger to the unprocessed forms (proIAPP_1-70_ K_*d*_ ~ 268±43 nM and proIAPP_1-51_ K_*d*_ ~ 329±29 nM) as compared to a relatively weak interaction with mature IAPP (K_*d*_ ~ 921±177 nM). The ability to discharge bound cargo in the acidic milieu of endosomes is a characteristic feature of endocytic receptors, including SORLA [13]. In line with this observation, binding of IAPP peptides to SORLA was strongest at pH 7.4, weaker at pH 5.5, and absent at pH 4.5 (Fig. 6d-f).

**Fig. 6.**
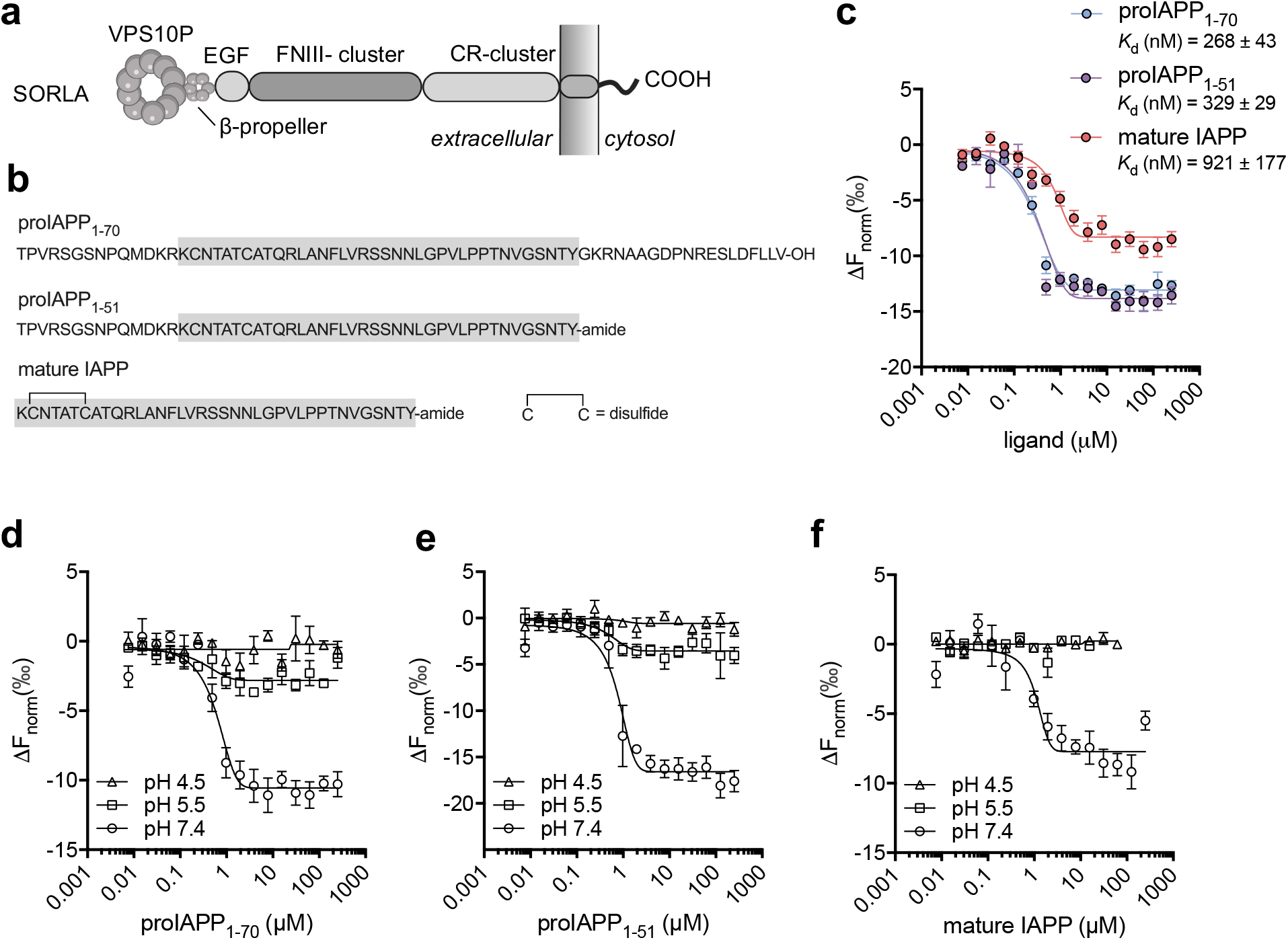
SORLA preferentially binds proIAPP in a pH-dependent manner. **(a)** Structural domains of SORLA, including vacuolar protein sorting 10 protein (VPS10P) domain, β-propeller, epidermal growth factor (EGF) repeats, clusters of complement-type repeats (CR), and fibronectin type III (FNIII) domains. **(b)** Amino acid sequences of murine proIAPP_1-70_, proIAPP_1-51_, and mature IAPP peptides. **(c)** Binding characteristics between the ectodomain of SORLA and murine IAPP peptides. The His-tagged ectodomain of human SORLA was recombinantly produced in HEK293-EBNA cells and purified using Ni^2+^ affinity chromatography as described in [12]. Binding of different forms of IAPP peptide to the ectodomain of SORLA was determined by microscale thermophoresis (MST) at pH 7.4. The concentration of the fluorescently-labeled SORLA ectodomain was kept constant (3 nmol/l), while non-labeled IAPP was serially titrated from 7.6 nmol/l – 250 μmol/l. Average *K_d_* was derived from at least 3 independent experiments. Data are shown as mean ± SEM. **(d-f)** Binding of (d) proIAPP_1-70_, (e) proIAPP_1-51_, and (f) mature IAPP to the SORLA ectodomain was tested by MST at pH 4.5, 5.5, and 7.4 as detailed in (c).

### SORLA mediates endocytosis of proIAPP, but not mature IAPP

Our studies indicate that SORLA may act as a receptor for IAPP peptides, possibly delivering them to lysosomal catabolism. To corroborate this hypothesis, we tested the ability of SORLA to mediate endocytic uptake of IAPP *in vitro*. Here, we assayed IAPP peptide uptake in a neuroblastoma SH-SY5Y cell line stably overexpressing SORLA. This cell line is commonly used to study SORLA-mediated sorting functions [12, 14]. Moreover, this cell line does not express endogenous IAPP (Fig. 7a, DMSO panel), enabling us to examine cellular uptake of unlabeled synthetic IAPP peptides.

**Fig. 7.**
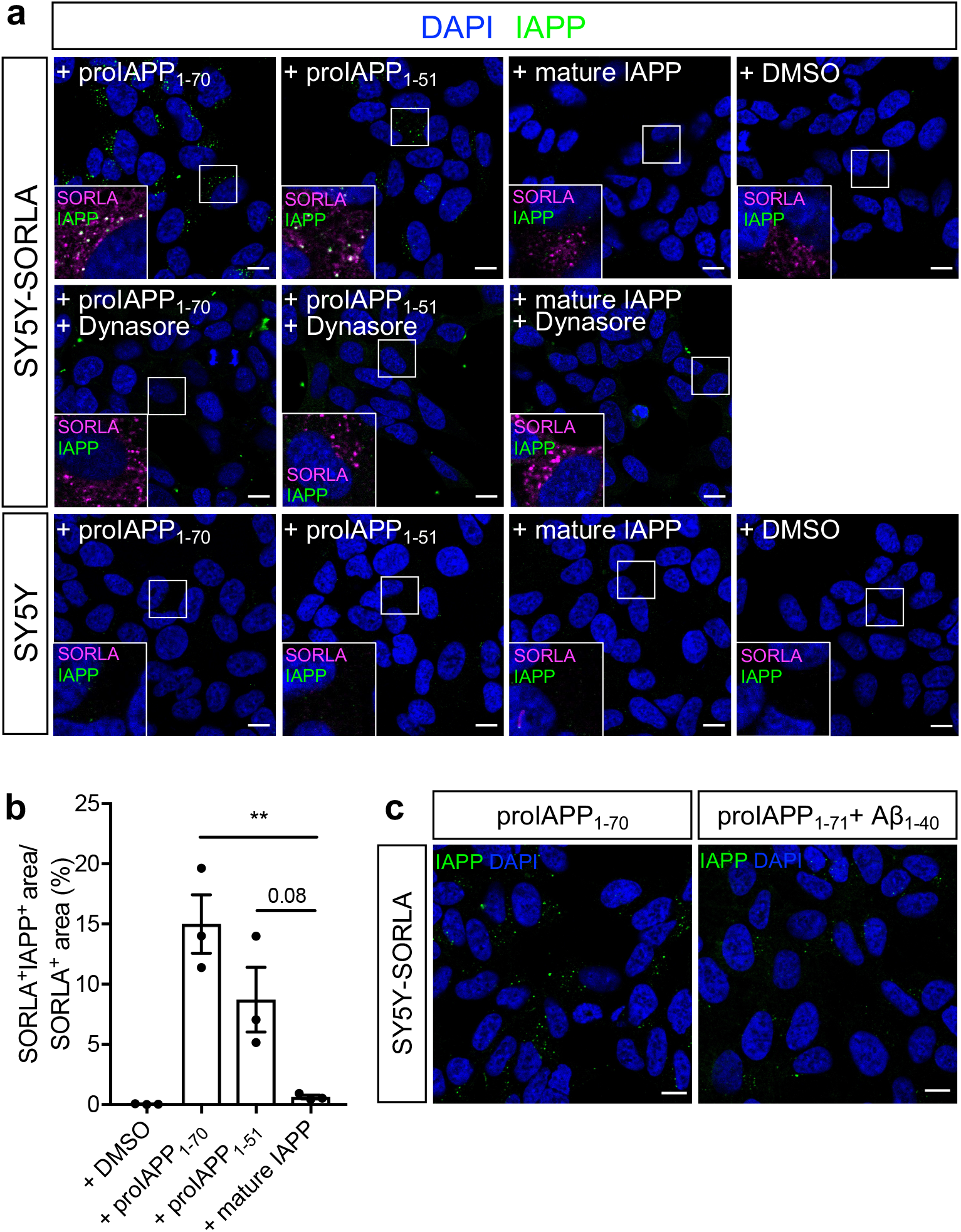
SORLA mediates endocytic uptake of proIAPP, but not mature IAPP. **(a)** SY5Y cells stably overexpressing SORLA (top, middle panels) and parental SY5Y cells (bottom panel) were incubated with 20 μmol/l of proIAPP_1-70_, proIAPP_1-51_, or mature IAPP peptides, or with 0.1% DMSO as solvent control. Where indicated, cells were also treated with 100 μmol/l dynasore to block dynamin-mediated endocytosis. After 30 minutes, the cells were fixed and immunostained for SORLA (magenta) and IAPP (green). Nuclei were counterstained with DAPI. The insets depict higher magnifications of the areas indicated by white boxes in the overview images. Scale bars, 10 μm. **(b)** The amount of internalized IAPP peptides was quantified based on the percentage of SORLA immunosignals colocalizing with each IAPP peptide (*n* =3 independent experiments, each with 3 – 4 images per condition). **(c)** Competition of proIAPP_1-70_ uptake in SY5Y-SORLA cells by Aβ_1-40_. Cells were treated with 20 μmol/l proIAPP_1-70_ alone or in the presence of 100 μmol/l Aβ_1-40_ for 30 minutes. Internalized proIAPP1-70 in cells were then immunolabeled by anti-IAPP antibody (green) and nuclei counterstained with DAPI (blue). Representative images from 3 independent experiments. Scale bars, 10 μm. Data are shown as mean ± SEM. Statistical significance of differences in (b) was determined by one-way ANOVA with post hoc test. ** p < 0.01.

When cellular uptake of exogenously added IAPP peptides in SY5Y cells expressing SORLA was evaluated using immunocytochemistry, intracellular accumulation was seen for proIAPP_1-70_ and proIAPP_1-51_, but not for mature IAPP (Fig. 7a, top panel). Further quantifications showed that proIAPP_1-70_ was the most readily internalized species in the presence of SORLA (Fig. 7b). SORLA-dependent uptake of the proIAPP peptides was inhibited by treatment with dynasore, an inhibitor of clathrin-mediated endocytosis (Fig. 7a, middle panel). No uptake was seen in parental SY5Y cells lacking SORLA expression (Fig. 7a, bottom panel).

Since SORLA is known to bind small peptides including Aβ via its VPS10P domain [21], we next tested if proIAPP_1-70_ also binds to the same region in the receptor’s ectodomain. We found that the level of internalized proIAPP_1-70_ in SORLA-expressing SY5Y cells was significantly reduced in the presence of a 5-fold molar excess of Aβ (Fig. 7c), indicating that proIAPP_1-70_ and Aβ compete for the same binding site in SORLA.

Finally, immunostaining revealed that internalized proIAPP peptides were predominately directed to early endosomes (EEA1, 89.9%) (Fig. 8). To identify whether endocytosed peptides were further transported to lysosomes for degradation or to the TGN for recycling, we stained with a dye for lysosomes (LysoTracker) or immunostained with TGN38, respectively. Our results showed that proIAPP peptides colocalize more with lysotracker (51.6%) than with TGN38 (25.2%) (Fig. 8), suggesting delivery to lysosomes rather than TGN recycling of internalized propeptides.

**Fig. 8.**
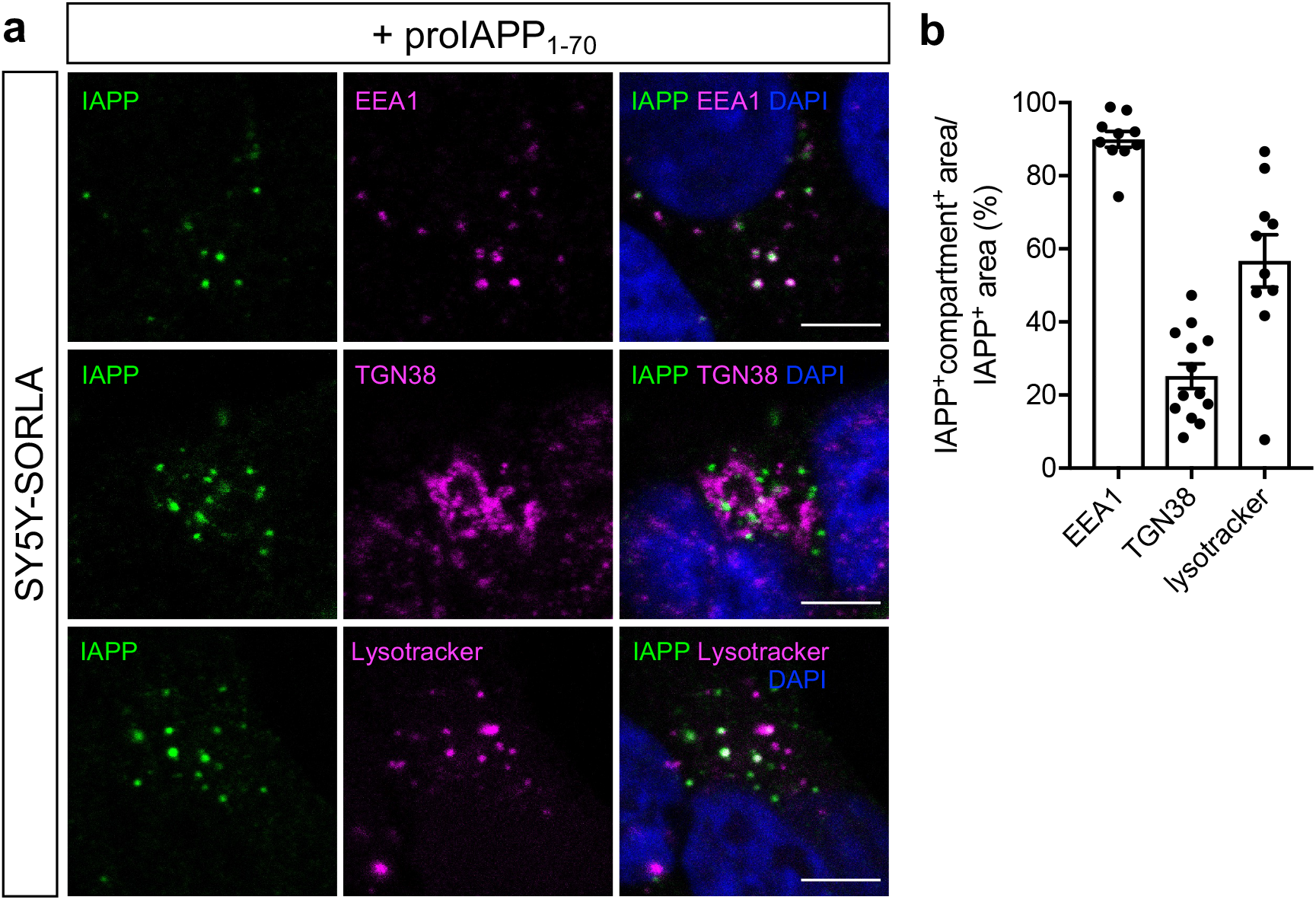
SORLA delivers proIAPP towards endolysosomal compartments. **(a)** SY5Y-SORLA cells were treated with 20 μmol/l proIAPP_1-70_ for 30 minutes as described in Fig. 7a. Subsequently, the cells were immunostained for IAPP (green) and compartment makers (magenta) EEA1, TGN38. Lysosomes were labeled by preincubating cells with Lysotracker Deep Red for 1 h prior to uptake assay. Scale bars, 5 μm. (**b**) Quantifications of object-based colocalization between internalized proIAPP_1-70_ and each compartment marker (*n* = 10-13 images per marker).

## Discussion

Increased levels of IAPP and its precursors due to overproduction or hypersecretion have been proposed as an underlying mechanism of islet amyloid formation. Therefore, maintaining IAPP homeostasis is essential to minimize the propensity for self-aggregation. We have identified a unique cellular pathway that counteracts this pathological potential of IAPP. In our model, SORLA acts as an endocytic receptor specific for the proform of the peptide released from islet beta cells. Receptor-mediated clearance of proIAPP delivers it to lysosomal catabolism, reducing extracellular buildup of this amyloidogenic peptide into fibrils and protecting islets from amyloid-induced cell death (Fig. 9).

**Fig. 9.**
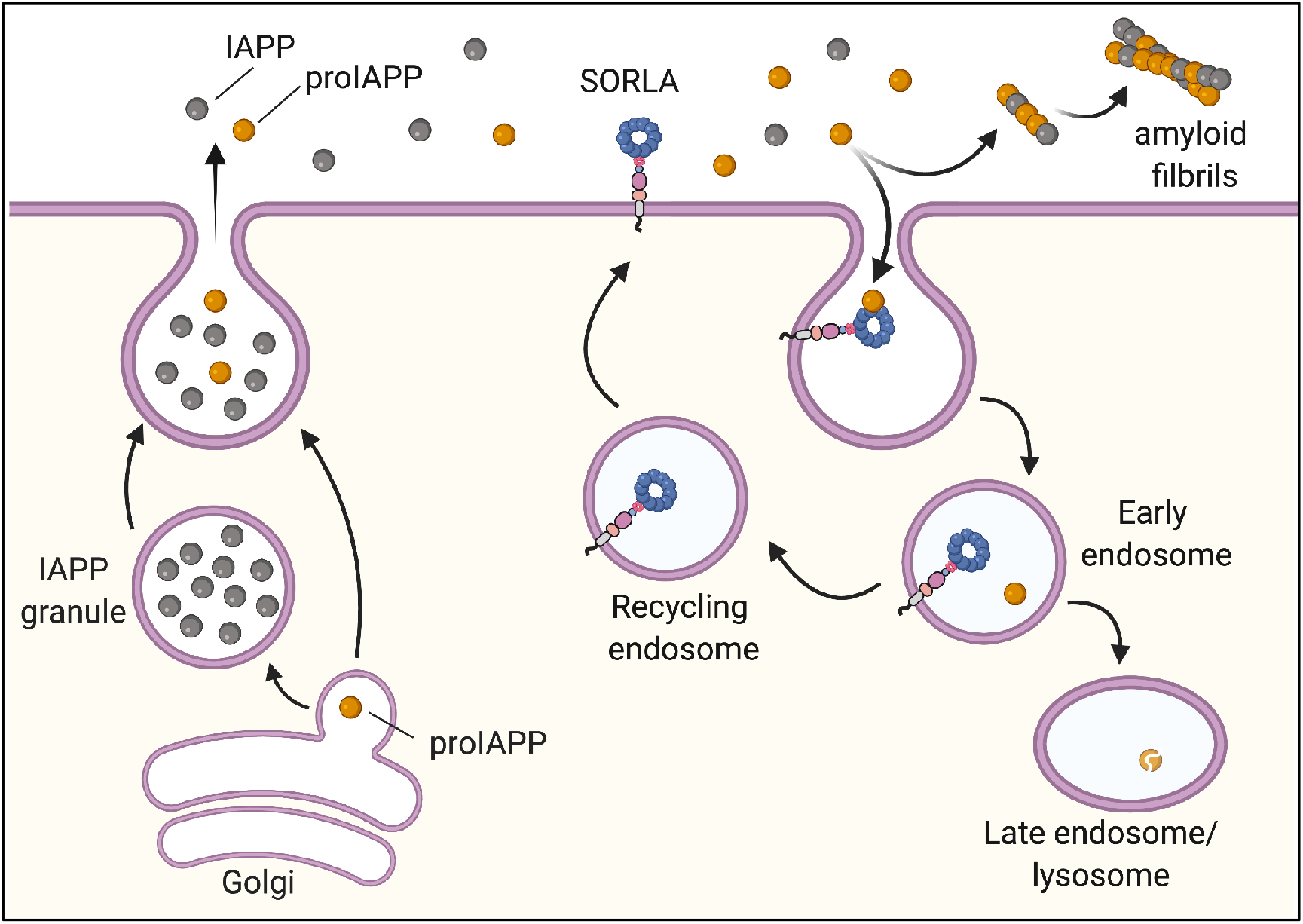
Proposed pathway of SORLA-mediated endocytosis of proIAPP. SORLA functions as an endocytic receptor for clearance of pro-but not mature IAPP released from islet beta cells. Following pH-dependent release of proIAPP in early endosomes, unliganded SORLA recycles back to the cell surface, whereas proIAPP is destined for lysosomal degradation. SORLA-mediated clearance of proIAPP released from cells counteracts pancreatic amyloid fibril formation. Created with BioRender.

The cellular pathways for IAPP biosynthesis are complex, involving movement of nascent proIAPP along the biosynthetic route from the endoplasmic reticulum to the Golgi as well as transport of its partially processed proIAPP intermediate to secretory granules for final processing and maturation. Extensive studies have identified prohormone convertases (PC1/3 and PC2) [28, 29], peptidyl-glycine alpha-amidating monooxygenase (PAM) [30] and carboxypeptidase E (CPE) [31] as key regulators of proIAPP processing. In addition to removing basic residues from propeptides, CPE is suggested to also function as a sorting receptor for directing prohormones including proinsulin to the regulated secretory granules [32]. However, conflicting results from other studies have challenged the role of CPE as a prohormone sorting receptor [33]. Following a hypothesis-driven approach, we explored a possible role for the VPS10P domain receptor SORLA in IAPP trafficking processes. VPS10P domain receptors are a distinct class of sorting receptors that direct intracellular trafficking and processing of proteins along the secretory and endocytic paths of cells [34]. With relevance to islet biology, the VPS10P domain receptor SorCS1 had been shown to direct replenishment of secretory granules with insulin [35], providing a molecular explanation for its association with type 2 diabetes in humans and mouse models [36, 37]. While our data dispute a similar function for SORLA in intracellular sorting and release of IAPP (Suppl. Fig. 3), they reveal a surprising role for this receptor in the clearance of extracellular proIAPP (Fig. 7). SORLA-mediated proIAPP clearance is blocked by dynasore, indicating a clathrin-dependent mechanism of uptake. In addition, internalized proIAPP molecules are preferentially sorted to endolysosomal compartments (Fig. 8), suggesting that they are destined for lysosomal catabolism. In support of this assumption, binding of IAPP to SORLA is lost at low pH (Fig. 6). pH dependency of binding is a hallmark of endocytosis, enabling receptors to discharge their cargo in acidified endosomes. Because of a low endocytic activity of islet beta cells *in vitro*, we have not been able to directly document SORLA-dependent uptake of proIAPP in this cell type. However, predominant localization of SORLA to early, late, and recycling endosomes in isolated beta cells (Fig. 5) suggests a similar endocytic receptor function as in SY5Y cells.

Our observation that SORLA-mediated uptake of proIAPP is impaired by Aβ (Fig. 7c) indicates that proIAPP binds to the same site as Aβ in a funnel formed by the VSP10P domain, a 700 amino acid module shared by the ectodomains of all VSP10P domain receptors [38]. Importantly, the affinity of SORLA is higher for the pro-than for the mature form of the peptide (K_*d*_ ~ 300 nM vs. 921 nM), explaining its ability to clear pro- but not mature IAPP. This observation is noteworthy as it addresses an open question concerning the origin of islet amyloid. Loss of the proIAPP receptor SORLA increases islet amyloid deposition in mice (Fig. 2). This finding supports earlier hypotheses that proIAPP is likely a critical species in initiating early steps of amyloid deposition [7, 39]. There have also been debates on the extracellular or intracellular origin of islet amyloid [40–42]. Our model of SORLA-mediated endocytosis of proIAPP provides new evidence to support islet amyloid formation in the extracellular space due to an aberrant accumulation of secreted proIAPP.

Significant amounts of islet amyloid are already present in normal chow-fed hIAPP-expressing mice lacking SORLA, but not in wildtype control. This finding is surprising as previous studies on hIAPP transgenic mice reported no spontaneous islet amyloidosis unless additional diabetogenic traits such as obesity or insulin resistance were introduced genetically [43], pharmacologically [22] or through dietary interventions [44]. Our data suggest that SORLA has an important role in maintaining IAPP homeostasis under normal physiological conditions. This protective effect against islet amyloid is not seen under conditions of dietary stress, possibly because it is masked by hypersecretions of hIAPP induced by HFD (Suppl. Fig. 3a-f).

The severity of islet amyloid deposition is correlated to beta cell dysfunction, apoptosis, and hyperglycemia [45–47]. *In vitro* studies have demonstrated that synthetic hIAPP-induced amyloid fibrils are toxic to beta cells in cultured islets [48], and that pharmacological inhibition of hIAPP fibrils formation improves viability of cultured islets [49]. In the present study, we observed aggravated cell death in islets of hIAPP-expressing SORLA-deficient mice *in vivo*, in line with enhanced islet amyloid in these animals. However, cell death did not progress to overt beta cell dysfunction and impairment in glucose metabolism under the experimental conditions tested here. Still, one may expect that the observed increase in cell death will have negative impacts on islet function in SORLA deficient mice at an older age, following further accumulate amyloid deposits.

In conclusion, our studies have identified SORLA-dependent catabolism of amyloidogenic peptides as a protective mechanism common to the brain and pancreas. Future studies will corroborate whether this receptor plays a similarly important role in glucose homeostasis and onset of diabetes as it does in neurodegeneration and risk of AD.

## Materials and methods

### Animal studies

Human IAPP transgenic (FVB/N-Tg(Ins2-IAPP)RHFSoel/J) mice were purchased from the Jackson Laboratory (#08232). SORLA deficient mice *(Sorl1^-/-^)* with a C57BL/6J background were previously generated [12]. SORLA WT and KO mice expressing hIAPP transgene and littermates hIAPP-null mice were generated by crossing hIAPP transgenic males with *Sorl1*^+/+^ or *Sorl1*^-/-^ females, respectively. All experiments were conducted in male mice. Animals were housed in a facility with controlled environment, 12-h light/ dark cycle, and fed a normal chow diet (4.5% crude fat) or a HFD (60% crude fat; #E15741-34; Ssniff, Germany). All experiments including measurements of body weight and fasting blood glucose, blood collection, intraperitoneal glucose tolerance test (GTT) and glucose-stimulated insulin secretion (GSIS) were performed according to protocols approved by the Berlin State Office for Health and Social Affairs (LAGESO, Berlin, Germany).

### Islet isolation, dispersion and culture

Animals were sacrificed by cervical dislocation and the pancreas was perfused with 2 ml of 900 U/ml collagenase (Sigma Aldrich, USA) in HBSS (Life Technologies, USA). Pancreas was removed and digested in 2 ml of collagenase solution at 37°C for 13 min, followed by manual shaking for 60 – 90 s, two rounds of washing and passed through a 70 μm filter. Islets were hand-picked and cultured in RPMI 1640 (PAN-Biotech, Germany), containing 11 mmol/l glucose, and supplemented with 2 mmol/l L-Glutamine, 100 U/mL penicillin, 100 mg/mL streptomycin and 10% FBS. Islets were recovered overnight prior to secretion assays or dispersion. For subcellular localization studies, islets were dispersed into single cells by pipetting in 0.05% trypsin-EDTA (Gibco, USA) solution for 1 min, seeded on uncoated glass coverslips and cultured for 6 days prior to fixation.

### Staining of tissues or cells for confocal microscopy

Pancreas tissues from mice were harvested and fixed in 4% (wt/vol.) paraformaldehyde overnight at 4°C, tissues were then infiltrated with sucrose in PBS stepwise from 15% to 30% (w/v), and preserved in freezing block containing in OCT compound. Tissue sections (10 μm) were rehydrated in 0.3% Triton X-100 in PBS-T for 15 min, followed by antigen retrieval in 10 mmol/l citrate buffer with 0.05% Tween-20 (pH 6.0) at 95°C for 10 min. For dispersed islet cells, cells were permeabilized in 0.3% Triton X-100, 0.1% BSA in PBS-T (pH 7.4) for 10 min at room temperature. Paraffin-embedded tissue sections from anonymized, healthy human pancreas biopsies were kindly provided by Prof. Søndergaard (Steno Diabetes Center Aarhus). Both tissues and cells were blocked in 3% BSA in PBS-T overnight at 4°C, followed by sequential incubation in primary and secondary antibodies (Suppl. Table 1) for 2 and 1 h, respectively at room temperature. Cell nuclei were visualized by DAPI staining. Images were acquired using a Zeiss LSM 700 confocal microscope (tissue sample with 10X objective; dispersed islet cells with 63X objective).

#### Amyloid staining

Islet amyloid was assessed based on thioflavin S (Sigma-Aldrich, USA) staining, as previously described [8]. Amount of islet amyloid deposition was quantified as thioflavin S positive area over total islet area in percentage, using CellProfiler (Cambridge, MA, USA).

#### TUNEL staining

Cell death was measured by terminal deoxynucleotidyl transferase dUTP nick end labeling (TUNEL) staining according to manufacturer’s protocol (Roche Applied Science, Germany).

### Dynamic insulin secretion assay by islet perifusion

Groups of 30 islets were used per sample and the assay was performed using a PERI 4.2 machine (Biorep Technologies, USA). Islets were continuously perifused with KRBH buffer (129 mmol/l NaCl, 4.8 mmol/l KCl, 1.2 mmol/l MgSO_4_, 1.2 mmol/l KH_2_PO_4_, 5 mmol/l NaHCO_3_, 2.5 mmol/l CaCl_2_, 10 mmol/l HEPES and 0.25% BSA, at pH 7.4) with the indicated glucose concentration at a flow rate of 100 μl/min. Islets were equilibrated in 11 mmol/l glucose KRBH for 60 min prior to stimulation in the following sequence: 11 mmol/l glucose (18 min), 1.67 mmol/l glucose (60 min), 16.7 mmol/l glucose (34 min), 1.67 mmol/l glucose (30 min), and 30 mmol/l KCl in 1.67 mmol/l glucose KRBH (10 min). At the end of assay, islets were collected and lysed in TE buffer (10 mmol/l Tris-HCl, 1 mmol/l EDTA and 1% Triton) to determine total cell DNA content (Quant-iT Picogreen DNA kit, Thermo Fisher Scientific, USA). Levels of secreted insulin were measured by ELISA and data were normalized to total DNA.

### ELISAs

Concentrations of insulin were measured using the Ultra Sensitive Mouse Insulin ELISA kit (Crystal Chem, USA). Human proIAPP_1-48_ and mature IAPP were measured by in-house ELISA as described [50].

### Microscale thermophoresis (MST)

Murine proIAPP_1-70_, proIAPP_1-51_ and mature amidated IAPP peptides (NCBI Reference Sequence: NP_034621) were commercially synthesized (Biosyntan, Germany). Peptides were dissolved in PBS and stored at −80°C prior to binding assays. Recombinant His-tagged SORLA ectodomain (including residue 728-1526) was previously purified for Surface Plasmon Resonance analysis as described in [12]. In MST, SORLA ectodomain was fluorescently labeled using the Protein Labeling Kit RED-NHS (NanoTemper Technologies, Germany). SORLA-labeled target molecule in PBS with 0.05 % Tween 20 (pH 7.4) was kept constant (3 nmol/l), while the concentration of the non-labeled binding ligand (IAPP) was serially titrated between 7.6 nmol/l – 250 μmol/l. *K_d_* was derived using MO.Affinity Analysis software version 2.3 (NanoTemper Technologies, Germany).

### Cell line and culture

Neuroblastoma SH-SY5Y cells (ATCC CRL-2266) were cultured in DMEM/F12 media (Gibco, USA) supplemented with 10% FBS, 1% NEAA, 100 U/mL penicillin and 100 mg/mL streptomycin. SH-SY5Y cells stably overexpressing SORLA were previously generated [14] and maintained in the presence of 90 μg/ml zeocin (Invitrogen, USA). Cells were routinely tested for mycoplasma infection.

### IAPP peptide uptake assay

The same synthetic murine (pro)IAPP peptides as described in MST were used in this assay. Synthetic human Aβ_1-40_ peptides were purchased from Bachem, Germany (#4095737). SH-SY5Y parental cells and SH-SY5Y cells stably overexpressing SORLA were seeded on glass coverslip one day prior to peptide uptake. Cells were incubated in serum-free medium for 30 min prior to treatment with 20 μmol/l (pro)IAPP for 30 min. Simultaneous treatment with 100 μmol/l dynasore (Cayman Chemical, USA) was used to examine the role of clathrin-mediated endocytosis. Cells were fixed in 4% (wt/vol.) paraformaldehyde and immunofluorescence staining was performed to visualize presence of internalized peptide, SORLA and subcellular organelles. Lysosomes were labeled by preincubating cells with 500 nmol/l LysoTracker Deep Red (Thermo Fisher Scientific, USA) in normal growth media for 1 h prior to uptake assay.

### Statistical analysis

Statistically analyses were performed using GraphPad Prism 6.0 (GraphPad Software, USA). Normally distributed datasets were analyzed using Student’s t-test, one-way or two-way ANOVA. Data are presented as mean ± SEM.

## Abbreviations

Aβ: Amyloid-beta peptide
APP: Amyloid precursor protein
HFD: High-fat diet
hIAPP: Human islet amyloid polypeptide
IAPP: Islet amyloid polypeptide
KO: Knockout
ND: Normal diet
SORLA: Sorting protein-related receptor containing LDLR class A repeats
WT: Wildtype

## Acknowledgement

We are indebted to C. Kruse and K. Kampf for expert technical assistance and to V. Schmidt-Krüger for help in obtaining animal experimentation licenses. We thank the Advanced Light Microscopy Platform and the Preclinical Research Center, of the MDC for technical support and assistance in this work. We thank Prof. Annette Schürmann and members of her group (German Institute of Human Nutrition, Potsdam-Rehbru□cke) for the use of, and assistance with the automated islet perifusion system. We also thank Prof. Esben Søndergaard (Steno Diabetes Center Aarhus, Aarhus) for providing tissue sections of human pancreas biopsies.

## Data availability

Data are available on request from the authors.

## Authors’ relationships and activities

The authors declare no competing interest related to this manuscript.

## Contribution statement

AZLS conceived and designed the study, performed experiments, collected and analyzed the data. YCC performed experiments, collected and analyzed data (human IAPP ELISA). CBV contributed to the design of experiments and provided critical reagents. TEW contributed to the design of experiments and interpreted data. AZLS and TEW wrote the manuscript with editorial input, review and approval for publication from all authors. AZLS and TEW are guarantors of this work.

