## Supplemental Methods for "SORLA mediates endocytic uptake of proIAPP and protects against islet amyloid deposition"

### Supplementary Methods

#### Human proIAPP$_{1\text{-}48}$ and IAPP secretions from isolated islets

Groups of 30 islets were collected in low-retention Eppendorf tubes and incubated in 200 µl of KRBH buffer with the indicated glucose concentration. Islets were first equilibrated in 11 mmol/l glucose KRBH for 1 h, followed by sequential 1 h stimulation with 11 mmol/l (culture) glucose, 1.67 mmol/l (low) glucose and 16.7 mmol/l (high) glucose. Islet supernatants were collected for measurement of human proIAPP$_{1\text{-}48}$ and mature IAPP by ELISA. Islets were washed and harvested for protein quantification. Levels of human proIAPP$_{1\text{-}48}$ and IAPP were normalized to total protein content.
