## Supplemental Table 1 for "SORLA mediates endocytic uptake of proIAPP and protects against islet amyloid deposition"

**Supplementary Table**

**Table 1. Information on antibodies for immunofluorescence staining**

| Antibody | Manufacturer | Catalog number | Host Species | Dilution |
|---|---|---|---|---|
| SORLA | In-house, Denmark | | Goat | 1:200 |
| Insulin | Agilent, USA | IR002 | Guinea pig | 1:3 |
| Mouse IAPP | Peninsula Laboratories, USA | T-4145 | Rabbit | 1:400 |
| Human IAPP | Peninsula Laboratories, USA | T-4149 | Rabbit | 1:400 |
| Glucagon | Abcam, USA | Ab10988 | Mouse | 1:500 |
| Somatostatin (SST) | Abcam, USA | Ab111912 | Rabbit | 1:250 |
| Pancreatic polypeptide (PPY) | Sigma-Aldrich, USA | AB939-I | Rabbit | 1:500 |
| Syntaxin6 (STX6) | BD bioscience, USA | BD610636 | Mouse | 1:200 |
| EEA1 | BD bioscience, USA | BD610457 | Mouse | 1:100 |
| Rab4 | Abcam, USA | Ab13252 | Rabbit | 1:100 |
| Rab4 | BD bioscience | BD610889 | Mouse | 1:100 |
| Rab9 | Invitrogen, USA | MA3-067 | Mouse | 1:100 |
| Rab11 | BD bioscience, USA | BD610657 | Mouse | 1:100 |
| TGN38 | BD bioscience, USA | BD610899 | Mouse | 1:50 |
| Anti-goat AlexaFluor488 | Abcam, USA | ab150129 | Donkey | 1:1000 |
| Anti-mouse AlexaFluor555 | Abcam, USA | Ab150106 | Donkey | 1:1000 |
| Anti-mouse AlexaFluor647 | Invitrogen, USA | A31571 | Donkey | 1:1000 |
| Anti-guinea pig Cy3 | Jackson ImmunoResearch, UK | 706-165-148 | Donkey | 1:1000 |
| Anti-guinea pig AlexaFluor647 | Merck Milipore | AP193SA6 | Donkey | 1:1000 |
| Anti-rabbit AlexaFluor555 | Invitrogen , USA | A31572 | Donkey | 1:1000 |
| Anti-rabbit AlexaFluor647 | Abcam, USA | Ab150075 | Donkey | 1:1000 |

All antibodies were validated by manufacturer
