## Supplemental Fig. 1 for "SORLA mediates endocytic uptake of proIAPP and protects against islet amyloid deposition"

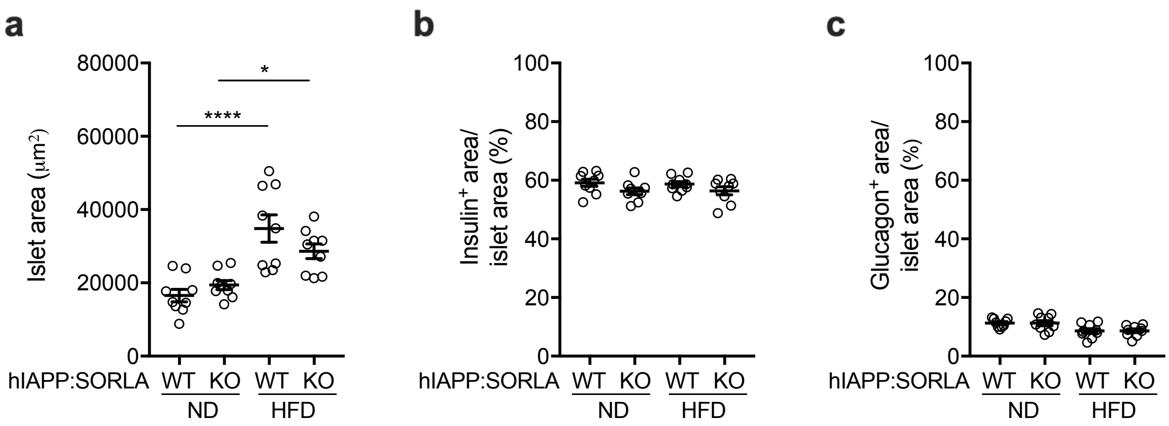


**Suppl. Fig. 1 SORLA deficiency does not impact islet size or islet cell composition in hIAPP-expressing mice.**

Quantifications of (**a**) total islet area or **(b)** beta cell (insulin$^{+}$) or (**c**) alpha cell (glucagon$^{+}$) area per islet area as determined by immunostainings of the respective markers on pancreatic sections from 33- to 35-weeks old mice of the indicated genotypes (immunostainings exemplified in Fig 3). Animals were fed a normal chow (ND) or a high fat diet (HFD) for 6 months ($n$ = 9 mice per genotype, 20 – 30 islets per mouse). Data are given as mean ± SEM. No statistically significant differences were detected between SORLA genotypes at the various experimental conditions. Statistical significance of differences in (a) was determined by two-way ANOVA with post-hoc test. $^{*}$ $p < 0.05$, **** $p < 0.0001$.
