## Supplemental Fig. 2 for "SORLA mediates endocytic uptake of proIAPP and protects against islet amyloid deposition"

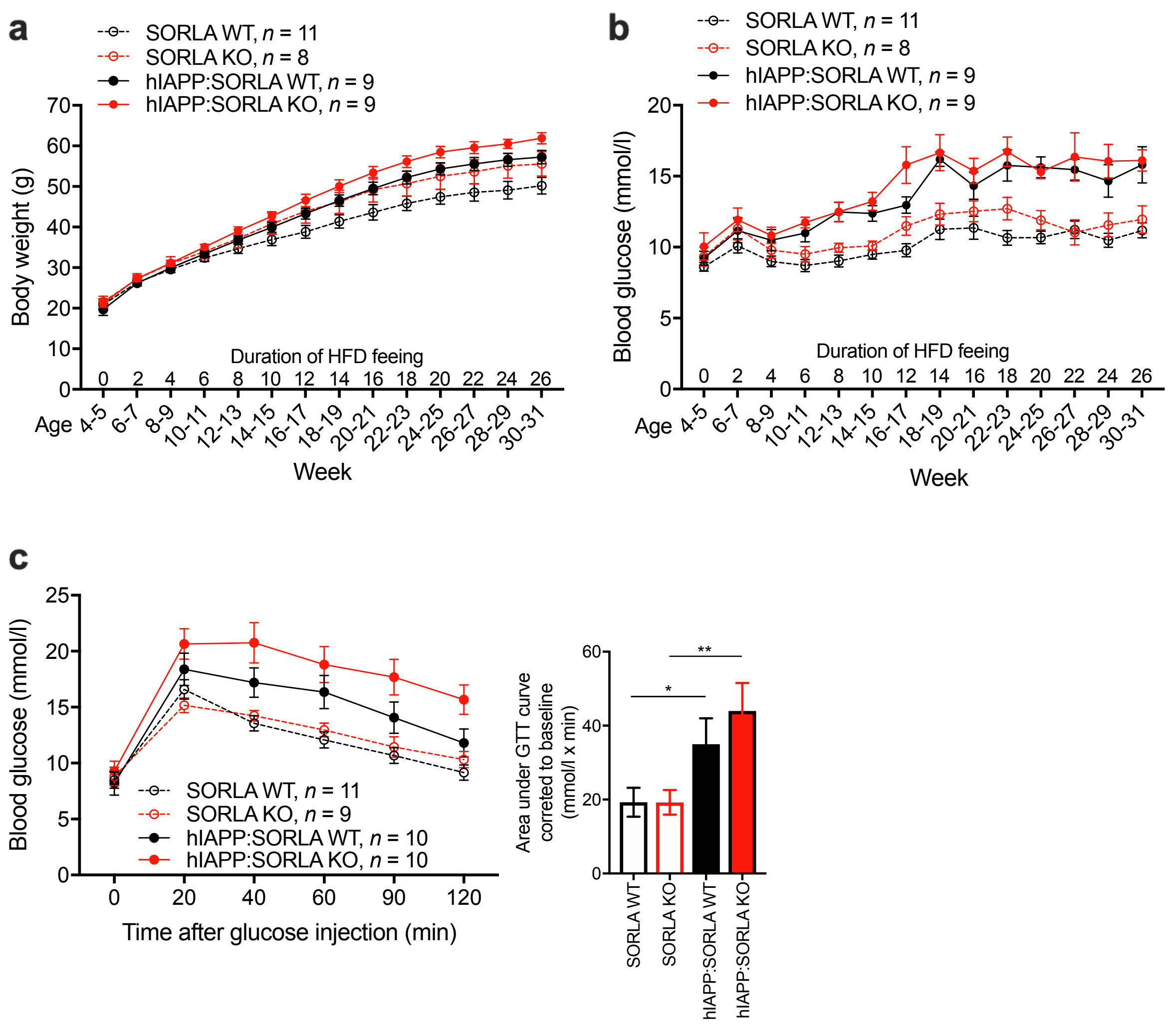


**Suppl. Fig. 2 Metabolic characterization of hIAPP-expressing wildtype and SORLA-deficient mice on a high fat diet.**

(**a, b**) Bi-weekly analysis of mice of the indicated genotypes for (a) body weight and (b) 6-h fasting blood glucose levels. **(c)** Glucose tolerance test (GTT) performed in 30- to 32-weeks old mice after a 16 h fast by intraperitoneal injection of 0.75 g/kg body weight of glucose. Response to glucose clearance was quantified based on the area under the GTT curves corrected to baseline. All data are expressed as mean ± SEM. Statistical significance of differences in (c) was determined by unpaired Student t-test. $^{*}$ $p$ < 0.05, ** $p$ < 0.01.
