## Supplemental Fig. 3 for "SORLA mediates endocytic uptake of proIAPP and protects against islet amyloid deposition"

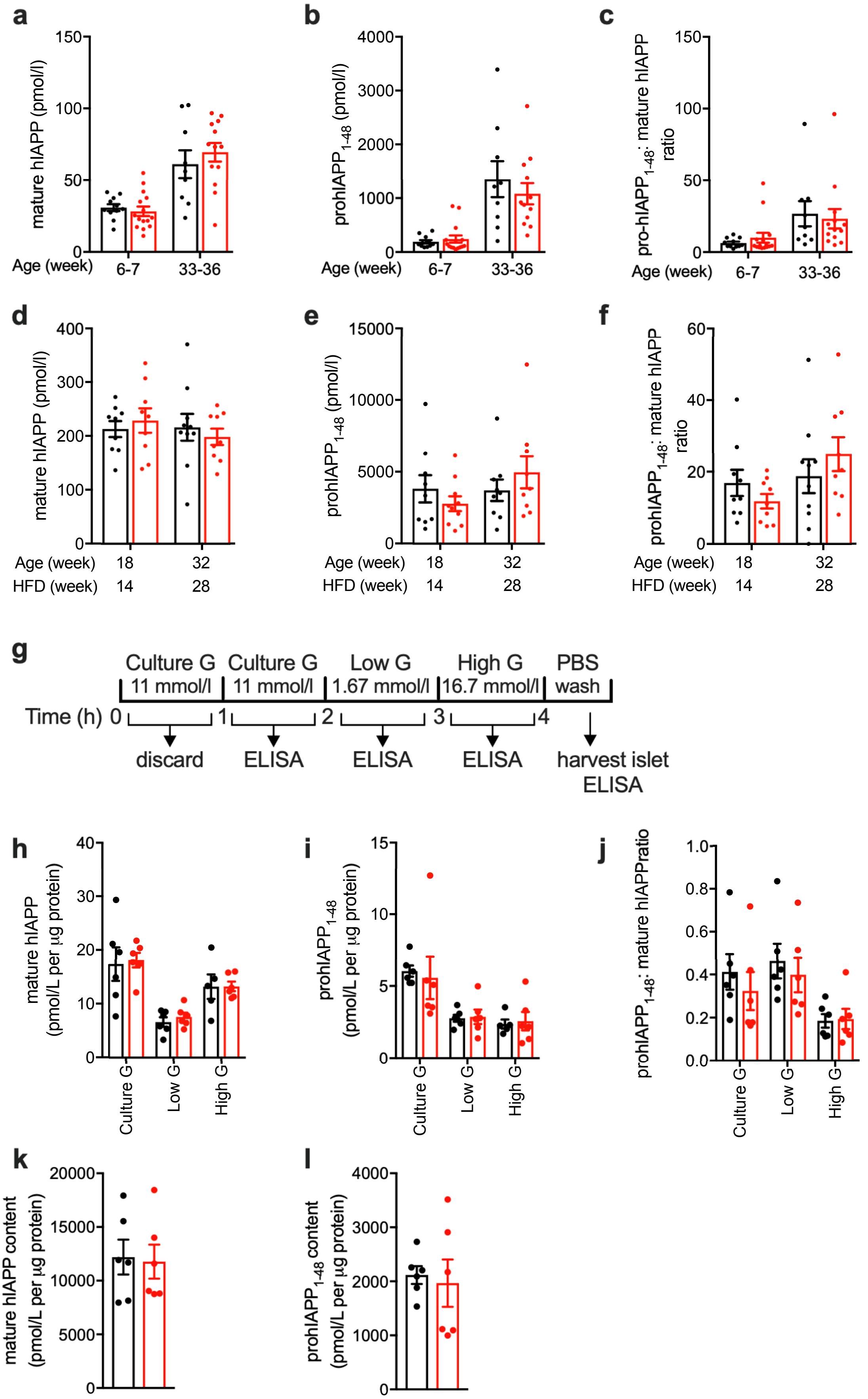


**Suppl. Fig. 3 SORLA deficiency does not impact hIAPP processing.**

(**a-f**) Fasting plasma levels of (a, d) mature hIAPP or (b, e) pro-hIAPP$_{1\text{-}48}$, as well as (c, f) pro- to mature-hIAPP ratio in hIAPP:SORLA WT (black bars) and hIAPP:SORLA KO (red bars) mice. Animals were fed a (a-c) ND or a (d-f) HFD. Analyses were performed at the indicated age (in weeks). No statistically significant differences in data were seen comparing genotypes using unpaired Student t-test (*n* = 6-15). (**g**) Schematic workflow of *in vitro* islet secretion assay. Islets were subjected to a series of incubations with secretion buffers containing the indicated concentrations of glucose (G). At the end of each incubation period, the buffer was collected for ELISA measurements. (**h-l**) Secreted levels of (h) mature hIAPP, (i) pro-hIAPP$_{1\text{-}48}$, (j) pro- to mature-hIAPP ratio as well as islet content of (**k**) mature hIAPP and (**l**) prohIAPP$_{1\text{-}48}$ in isolated islets of ND-fed hIAPP:SORLA WT (black bars) and hIAPP:SORLA KO (red bars) mice. Data are expressed as mean ± SEM. No statistically significant differences in data were seen comparing genotypes by unpaired Student t-test (*n* = 6).
